# ATR repression at telomeres by POT1a and POT1b: RPA exclusion and interference by CST

**DOI:** 10.1101/354985

**Authors:** Katja Kratz, Titia de Lange

**Affiliations:** Laboratory for Cell Biology and Genetics, Rockefeller University; New York, New York, 10021; USA

**Keywords:** telomere, ATR signaling, RPA, single-stranded DNA, POT1a, POT1b, 3’ overhang, CST, Ctc1, Stn1, Ten1, Polymeraseα/primase

## Abstract

Telomeres carry a constitutive 3’ overhang that can bind RPA and activate ATR signaling. POT1a, a single-stranded (ss) DNA binding protein in mouse shelterin, has been proposed to repress ATR signaling by preventing RPA binding. Repression of ATR at telomeres requires the TPP1/TIN2 mediated tethering of POT1a to the the rest of the shelterin complex situated on the ds telomeric DNA. The simplest version of the tethered exclusion model for ATR repression suggests that the only critical features of POT1a are its connection to shelterin and its binding to ss telomeric DNA binding. In agreement with the model, we show that a shelterin-tethered RPA70 mutant, lacking the ATR recruitment domain, is effective in repressing ATR signaling at telomeres. However, arguing against the simple tethered exclusion model, the nearly identical POT1b subunit of shelterin is much less proficient in ATR repression than POT1a. We now show that POT1b has the intrinsic ability to fully repress ATR but is prevented from doing so when bound to the CST/Polα/primase complex. The data establish that shelterin represses ATR with a tethered ssDNA-binding domain that excludes RPA from the 3’ overhang and suggest that ATR repression does not require the interaction of POT1 with the 3’ end or G4 DNA.

## INTRODUCTION

One of the many tasks of the telomeric shelterin complex is to repress the ATR-dependent DNA damage response at the natural ends of chromosomes (reviewed in (Lazzerini-Denchi and Sfeir, 2016; Palm and de Lange, 2008)). The main mode of ATR activation involves the binding of the abundant trimeric RPA (RPA70, RPA32, and RPA14) complex to ssDNA, recruitment of ATR through an interaction between RPA70 and the ATR interacting partner ATRIP, loading of the 9-1-1 (Rad9-Hus1-Rad1) complex on the end of the duplex, and activation of ATR by 9-1-1-bound TopBP1 (reviewed in (Ciccia and Elledge, 2010)). A second pathway involves activation of ATR by RPA-bound ETAA1 and is independent of 9-1-1 and TopBP1 (Bass et al., 2016; Haahr et al., 2016).

The two DNA structures required for TopBP1-dependent ATR activation are present at telomeres, which carry a 9-1-1 loading site and a 3’ overhang of TTAGGG repeats that can bind multiple RPA trimers. Although telomeres often occur in the t-loop configuration (Griffith et al., 1999; Doksani et al., 2013), this structure is unlikely to guard against ATR signaling since the base of the t-loop retains the critical features required for loading of RPA and 9-1-1. Rather, ATR repression is achieved by the POT1 subunit of shelterin, which bind to the ss TTAGGG repeats (Denchi and de Lange, 2007). Deletion of POT1 leads to activation of ATR at most telomeres in a manner that is dependent on RPA and TopBP1 (Gong and de Lange, 2010). Superresolution STORM imaging showed that this ATR activation takes place despite the presence of t-loops (Doksani et al., 2013).

POT1 proteins bind the sequence 5’ TTAGGGTTAG 3’ either at a DNA end or at an internal position using two oligosaccharide/oligonucleotide binding (OB) folds in their N-terminal half (Loayza et al., 2004; Lei et al., 2004; Baumann and Cech, 2001). The interaction of POT1 with ss TTAGGG repeats is neither sufficient nor necessary to target the protein to telomeres (Loayza and de Lange, 2003; Liu et al., 2004; Ye et al., 2004; Kibe et al., 2010). Accumulation of POT1 at telomeres requires its association with shelterin, which is mediated by the binding of TPP1 to the C-terminal half of POT1 (Liu et al., 2004; Ye et al., 2004; Rice et al., 2017). TPP1 binds to TIN2 in shelterin, which in turn interacts with the duplex telomeric DNA binding proteins TRF1 and TRF2, thus anchoring POT1 on the duplex telomeric DNA.

As a result of a gene duplication event, rodent telomeres contain two closely related POT1 proteins, POT1a and POT1b, which have diverged in function. Both POT1a and POT1b can repress ATR kinase signaling in G1 but repression of ATR in S/G2 requires POT1a (Palm et al., 2009; Hockemeyer et al., 2005; Gong and de Lange, 2010). In S phase, POT1b, but not POT1a, governs the post-replicative generation of the telomeric 3’ overhang (Wu et al., 2012; Hockemeyer et al., 2006; Hockemeyer et al., 2008). In part, POT1b executes this function through the recruitment of CST (Ctc1, Stn1, Ten1; reviewed in (Price et al., 2010)), an RPA-like complex that binds ssDNA and interacts with Polymerase α/primase (Polα/primase) to mediate fill-in synthesis that restores the correct overhang length.

Telomeres lacking POT1 proteins accumulate RPA, which is not detectable at functional telomeres (Gong and de Lange, 2010). This observation suggested a simple competition model whereby the binding of POT1 to the ssDNA prevents RPA-dependent ATR activation. As RPA is ~200 fold more abundant than POT1a and POT1b and has the same sub-nanomolar affinity for ss telomeric DNA (Takai et al., 2011), it is unlikely that simple competition could explain the repression of ATR. Indeed, POT1 is not able to compete with RPA for ss telomeric DNA in vitro (Flynn et al., 2011). It was therefore proposed that the repression of ATR requires the local tethering of the POT1 proteins to the rest of shelterin, thus increasing the local concentration of POT1. Consistent with this tethering model, repression of ATR requires the association of the POT1 proteins with TPP1 and the TPP1- and TIN2-mediated interaction with TRF1 and/or TRF2. ATR activation throughout the cell cycle is observed when telomeres lack either TPP1 or TIN2 (Kibe et al., 2010; Takai et al., 2011), or when telomeres contain alleles of TPP1 or TIN2 that do not recruit POT 1a/b (Frescas and de Lange, 2014).

In its simplest form, the RPA exclusion model predicts that any shelterin-tethered protein with the ability to bind ss TTAGGG repeats can repress ATR signaling at telomeres. However, this prediction is not met in the context of POT1b, which can repress ATR signaling in G1 but not in S phase. POT1b and POT1a are equally abundant at telomeres, interact with TPP1 in the same way, and have indistinguishable DNA binding features (Hockemeyer et al., 2007; Palm et al., 2009; Kibe et al., 2010; Takai et al., 2011). Thus, the current knowledge of POT1b argues against the idea that any shelterin-tethered ss TTAGGG repeat binding protein can exclude RPA from telomeres and thus repress ATR signaling.

There are alternative models for how POT1 represses ATR signaling. The ability of human POT1 to protect telomeric G4 DNA from unfolding by RPA has been proposed to repress ATR signaling (Ray et al., 2014). It is also possible that the greater affinity of POT1 for the telomeric 3’ end (when ending in TTAG-3’ (Lei et al., 2004)) protects telomeres by providing POT1 with a competitive advantage over RPA. In another proposal, an S/G2-specific interplay between hnRNPA1, an RNA binding protein with high affinity for ss TTAGGG repeats, and the telomeric lncRNA TERRA serves an intermediary function in removing RPA from telomeric DNA and then allowing POT1 to bind (Flynn et al., 2011). Finally, it is possible that the POT1 proteins interfere with 9-1-1 loading. Although POT1a/b show not recognize the telomeric ds-ss junction in vitro (Palm et al., 2009), it is not excluded that the junction is bound by POT1 in the context of the whole shelterin complex.

Here we report that, consistent with the tethering model, ATR signaling can be partially repressed by the DNA binding domain of RPA70 when it is tethered to shelterin. Furthermore, we solve the POT1b conundrum by showing that POT1b, like POT1a, can repress ATR signaling in S phase but that its interaction with CST prevents it from doing so. The results provide evidence for the idea that ATR repression involves RPA exclusion by a shelterin-tethered ssDNA binding protein with no other specialized features.

## RESULTS

### Engineering a synthetic shelterin-tethered ATR repressor

We sought to test the idea that the exclusion of RPA by POT1 is based on the tethering of the ssDNA binding domain of POT1 to the duplex telomeric DNA rather than POT1 specific DNA binding features (e.g. binding to G4 DNA, binding to the 3’ end, or binding to junctions) (Figure 1A). We argued that the most stringent test of this tethering model would involve creating an RPA-based synthetic ATR repressor. RPA has the same affinity for ss TTAGGG repeats as POT1 (Takai et al., 2011) but a different DNA binding mode: Whereas POT1 binds DNA with two OB-folds, RPA uses three OB-folds (OB-A, -B, and -C) in RPA70 (Figure 1B) and one OB-fold in RPA32 (Fan and Pavletich, 2012). The C-terminal OB-fold (OB-C) of RPA70 interacts with RPA32, which in turn binds to RPA14.

**Figure 1.**
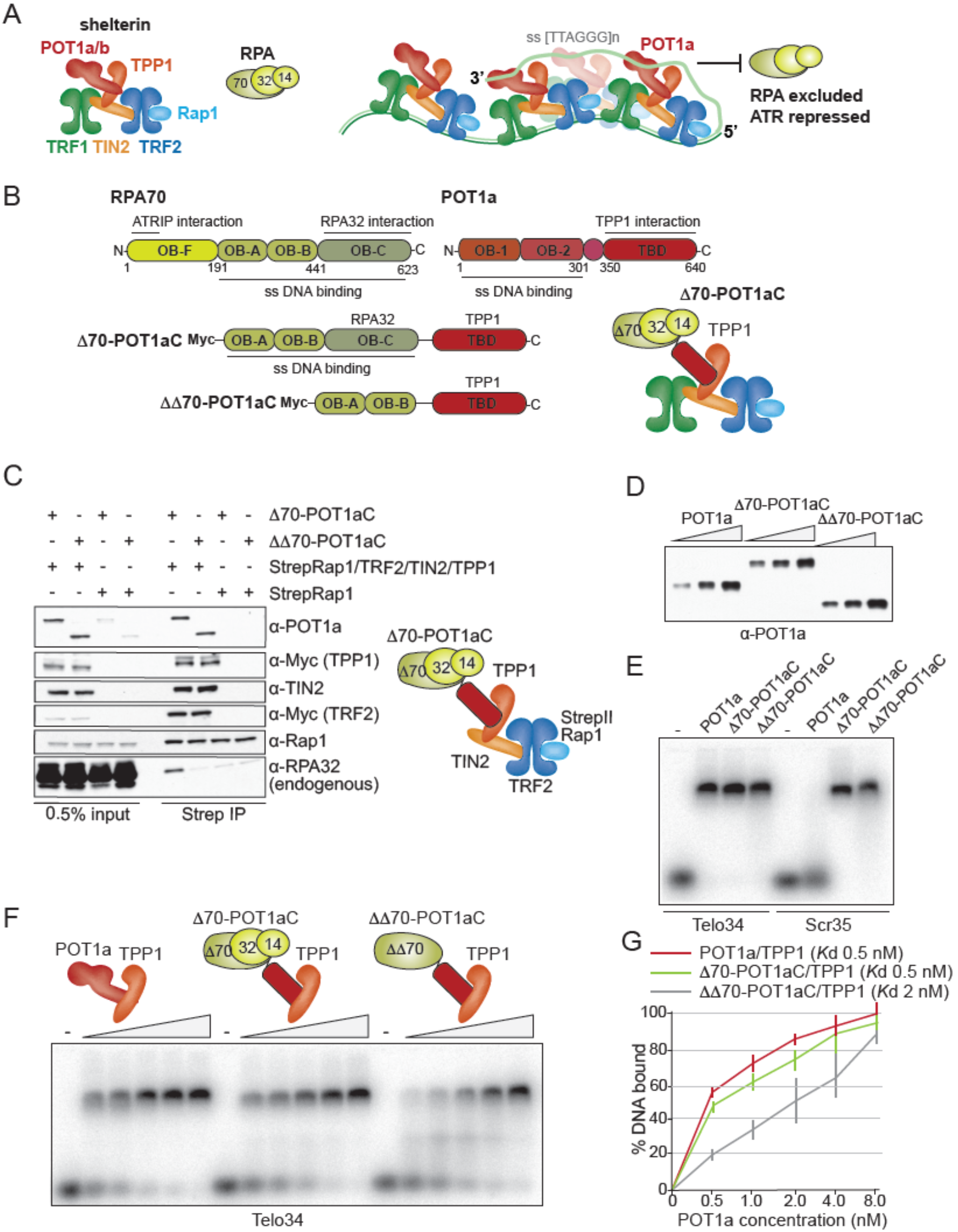
Characterization of the RPA-POTIaC fusion proteins. (A) Schematic (left) of the mouse shelterin complex and RPA and model (right) for how the binding of POT1a to shelterin and ss TTAGGG repeats excludes RPA from telomeres and thereby prevents ATR activation. (B) Schematic of mouse RPA70, mouse POT1a and the two fusion proteins Δ70-POT1aC and ΔΔ70-POT1aC, indicating the amino acids and functions of the relevant domains. Right: Schematic of mouse TPP1-tethered Δ70-POT1 aC and RPA32 and RPA14. (C) Immunoblots showing co-IPs from transiently transfected 293T cells of Δ70-POT1aC or ΔΔ70-POT1aC with either Strep-tagged Rap1 or Strep-tagged Rap1 together with TRF2, TIN2 and TPP1. Antibodies to the expressed proteins and the endogenous RPA32 are indicated. (D) Example of immunoblotting to determine the relative protein concentration of Strep-tagged TPP1/POT1a or TPP1/RPA-fusion proteins isolated from transfected 293T cells. (E) EMSA showing binding of TPP1/POT1a, TPP1/Δ70-POT1aC and TPP1/ΔΔ70-POT1aC to telomeric and non-telomeric single-stranded DNA. Telo34: 34 nts of single-stranded telomeric DNA; Scr35: 35 nts of single-stranded non-telomeric DNA (see Experimental Procedures for probe sequences). (F)Representative EMSAs to determine the apparent relative K_d_s of TPP1/POT1a, TPP1/Δ70-POT1aC and TPP1/ΔΔ70-POT1aC for Telo34.

An obvious complication in creating an RPA-based ATR repressor is that tethering of wild type RPA to telomeres would permanently recruit and likely active the ATR kinase. The interaction of RPA70 with the ATRIP/ATR complex is mediated by its fourth (N-terminal) OB-fold (OB-F), which does not bind to DNA (Xu et al., 2008). We therefore based the chimera on a version of RPA70 that lacks OB-F (referred to as Δ70) (Figure 1B). This part of RPA70 was fused to the C-terminus of POT1a (referred to as POT1aC), which interacts with TPP1 and ensures association with shelterin bound to duplex telomeric DNA.

To test for interaction with shelterin, Δ70-POT1aC and a second chimera, ΔΔ70-POT1aC, which also lacks the C-terminal RPA70 OB-fold, were co-expressed with TPP1, TIN2, TRF2, and StrepII-tagged Rap1 in 293T cells (Figure 1C). Immunoblotting of proteins associated with StrepII-Rap1 showed that both chimeras had the ability to form a complex with TPP1/TIN2/TRF2/Rap1. As expected, Δ70-POT1aC showed an interaction with RPA32, whereas ΔΔ70-POT1aC did not.

To determine the DNA binding features of the chimeric proteins, they were coexpressed with StrepII-tagged TPP1 (Figure 1D-G; Supporting Figure S1). The concentration of the isolated heterodimers was adjusted based on immunoblotting (Figure 1D) with an antibody to the C-terminus of POT1a and the complexes were analyzed in a gel-shift assay with labeled 34 or 35 nt probes (Figure 1E-G). The chimeric heterodimers formed with TPP1 bound to ss TTAGGG repeats as well as a scrambled sequence probe whereas the POT1a/TPP1 heterodimer only bound telomeric DNA (Figure 1E). Titration experiments showed that Δ70-POT1aC/TPP1 had the same affinity (apparent relative K_d_ ~0.5 nM) for TTAGGG repeats as POT1a/TPP1 (Figure 1F and G). Consistent with the absence of the third RPA70 OB-fold, the ΔΔ70-POT1aC/TPP1 heterodimer showed a lower affinity (apparent relative K_d_ ~2 nM) for ss TTAGGG repeats.

### Repression of ATR by Δ70-POT1aC

To test the ability of the chimeric proteins to prevent the activation of ATR at telomeres (Figure 2A), they were expressed in SV40-LT immortalized POT1a^F/F^ MEFs from which POT1a can be deleted with the Cre recombinase (Hockemeyer et al., 2006) (Figure 2B). As controls, the POT1aC fragment and the Δ70 mutant of RPA70 were expressed individually. Expression vectors were chosen to yield moderate expression levels of the chimeric proteins, because an excess of the mutant forms of RPA70 (Δ70 and ΔΔ70) might interfere with the essential functions of RPA in DNA replication and ATR signaling. Immunoblotting with RPA70 and POT1a antibodies showed that while all exogenous proteins were overexpressed compared to the endogenous POT1a, their abundance was far below the endogenous RPA70 (Figure 2B), making it unlikely that that the RPA mutant proteins would have a deleterious effect. Indeed, cells expressing the mutant RPA70 alleles showed no overt proliferation defect and had comparable S phase indices (Supporting Figure S2A and S2B). As expected, all exogenous proteins that carried the C-terminus of POT1a localized to telomeres (Figure 2C and D).

**Figure 2.**
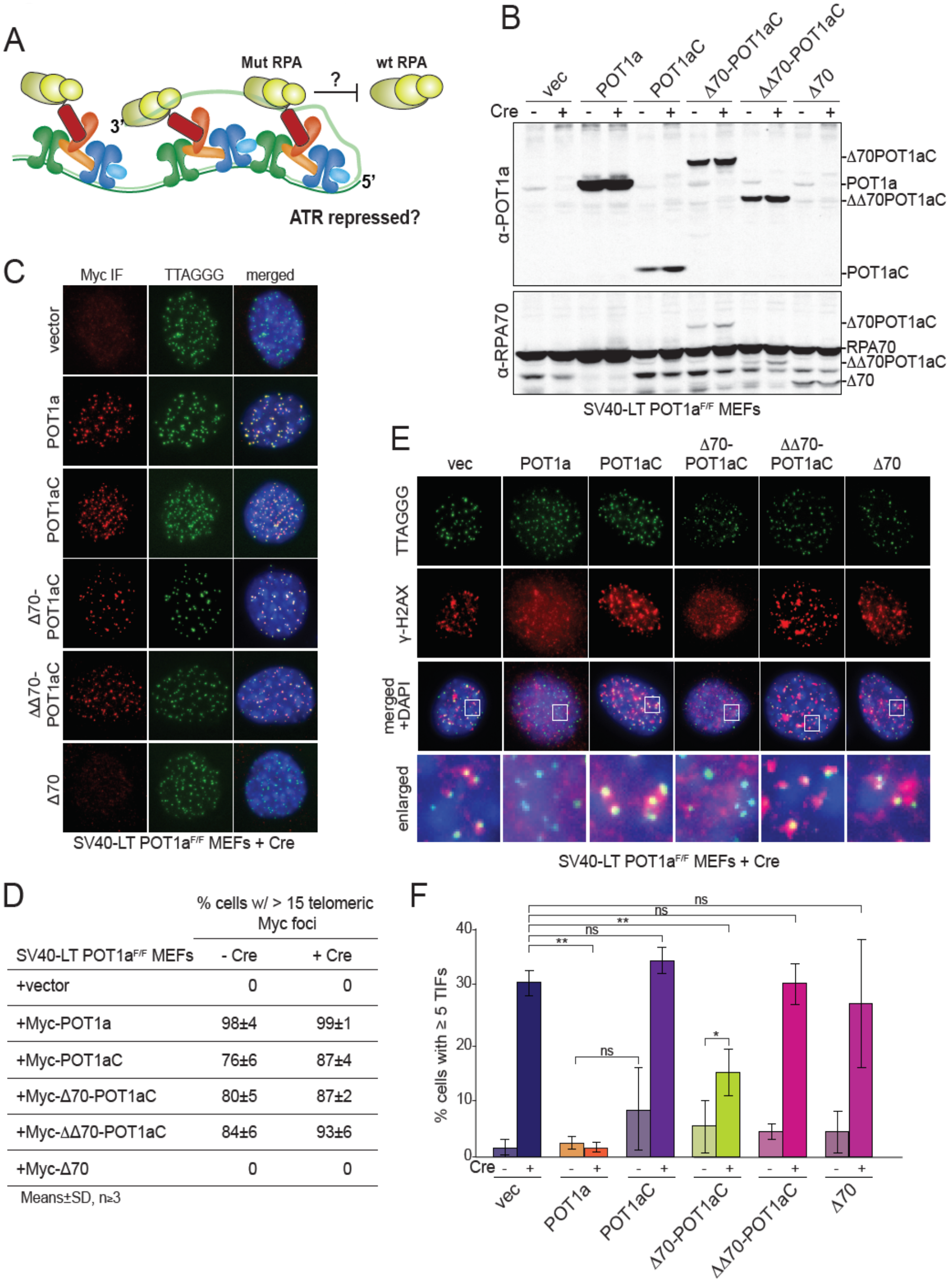
A shelterin-tethered RPA70 mutant can repress ATR signaling. (A) Schematic of the experimental set up. Shelterin-tethered Δ70-POT1 aC or (ΔΔ70-POT1 aC) are tested for their ability to replace the role of POT 1 a in ATR repression. (B) Immunoblot for POT1a and RPA70 in SV40-LT immortalized POT1a^F/F^ MEFs expressing the indicated proteins before and after Cre treatment (96 h time point). The endogenous POT1a is lost upon Cre treatment and the introduced proteins are overexpressed compared to endogenous POT1a (top). In contrast, the expressed RPA70 proteins are much less abundant than the endogenous RPA70 (bottom). The bands running below RPA70 are assumed to be degradation products. (C) Representative IF images showing the localization of the indicated exogenous Myc-tagged proteins in POT1a^F/F^ cells after deletion of the endogenous POT1a with Cre (96 h). IF was performed with an α-Myc antibody (red) together with FISH for telomeric DNA (green). (D) Quantification of the telomeric localization (as in panel C) of the indicated proteins in POT1a^F/F^ MEFs with and without Cre treatment (96 h). Averages of 3 independent experiments and SDs. (E) Representative IF images of the DNA damage response at telomeres of Cre-treated POT1a^F/F^ cells expressing the indicated proteins (96 h). γH2AX foci that co-localize with telomeres (TIFs) were detected with α-γH2AX antibody (red) combined with FISH for telomeric DNA (green). (F) Quantification of the TIF response as shown in (E) in the indicated POT1a^F/F^ cells with and without Cre treatment. Data represent averages from 3-5 independent with SDs. p-values were based on a two-tailed students t-test. *** ≤ 0.001; ** ≤ 0.01; * ≤ 0.05; n.s.: not significant.

Deletion of POT1a showed the expected induction of a DNA damage response at telomeres. In absence of POT1a, approximately 30% of the cells showed accumulation of γ-H2AX and 53BP1 at telomeres (Figure 2E and F; Supporting Figure S2C and D). This percentage of cells showing a DNA damage response is consistent with prior reports that showed that POT1a is required for the repression in S/G2 whereas in G1, POT1a is redundant with POT1b (Hockemeyer et al., 2006; Gong and de Lange, 2010).

As expected, this S/G2 ATR signaling was fully repressed by complementation of the cells with wild type POT1a, but not by POT1aC (Figure 2E and F). Remarkably, Δ70-POT1aC showed a significant reduction in ATR signaling at telomeres, resulting in only 15% of cells with detectable γ-H2AX foci at their telomeres. As expected, the untethered Δ70 protein was unable to repress ATR signaling at telomeres (Figure 2E and F). Cell cycle analysis indicated that the changes in ATR signaling were not due to changes in S phase index (Supporting Figure S2A and S2B) and quantitative analysis of the ss TTAGGG repeats at telomeres (Supporting Figure S2E) showed that the chimeric protein did not reduce the telomeric overhang signal, which could have confounded the interpretation of the reduced ATR signaling. In fact, overexpression of both POT1a and Δ70-POT1aC induced a slight increase in the telomeric overhang signal, presumably because they compete with POT1b for TPP1 interaction.

In contrast to Δ70-POT1aC, ΔΔ70-POT1aC was incapable of blocking ATR signaling at telomeres. We also tested the ability of E. coli SSB to repress ATR signaling when tethered to shelterin (Supporting Figure S3). The SSB-POT1aC chimera, which binds telomeric DNA with an apparent relative K_d_ of ~4 nM (Supporting Figure S3A-D), localized to telomeres but did not repress the DNA damage response (Supporting Figure S3E-H). We do not know whether the higher relative K_d_ of ΔΔ70-POT1aC and SSB-POT1aC is the cause of the lack of protection by these chimeras. In principle, in the context of tethered chimeras, changes in the relative K_d_ should not have a great impact. Possibly, the chimeras are defective in other aspects (e.g. exact positioning of the ssDNA binding domains within the shelterin complex).

### CST binding interferes with the repression of ATR by POT1b

It remains to be explained why POT1b is incapable of repressing ATR in S/G2 while it can do so in G1. POT1b interacts with CST, allowing appropriate fill-in synthesis of the telomere terminus during S/G2 (Wu et al., 2012). We therefore asked whether the interaction of POT1b with CST interferes with its ability to repress ATR. Prior work had shown that a POT1b allele with mutations in two sites of the protein (Figure 3A) does not bind to CST and is deficient in regulating the post-replicative processing of the telomere terminus (Wu et al., 2012). Notably, expression of this mutant version of POT1b fully restored the ATR repression in POT1a-deficient cells. The wild type POT1b, even when overexpressed, did not have this effect (Figure 3B-D).

**Figure 3.**
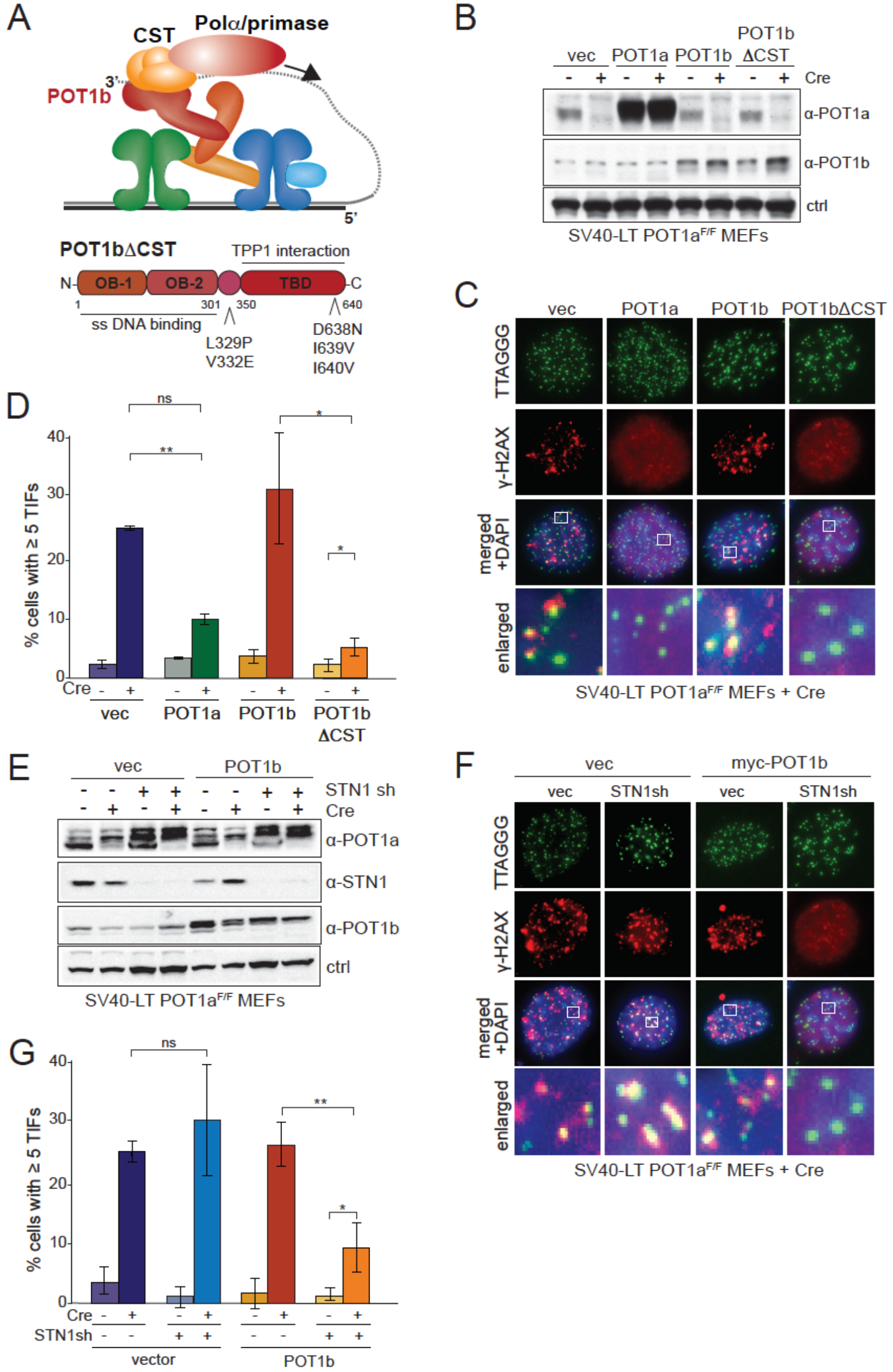
ATR repression by POT1b is limited by its interaction with CST. (A) Schematic of the role of POT1b in recruiting CST and Polymerasα/primase to telomeres (top). Schematic of POT1b domains and the mutations that disrupt the POT1b-CST interaction (bottom). (B) Immunoblot for POT1b in POT1aF/F MEFs expressing the indicated proteins before and after treatment with Cre (96 h). Ctrl: non-specific band used as loading control. (C) Representative IF-FISH images showing the DNA damage response at telomeres in Cre-treated POT1a^F/F^ MEFs expressing POT1a, POT1b, or POT1bΔCST (96 h). γH2AX foci that co-localize with telomeres (TIFs) were detected using α-γH2AX antibody (red) and a telomeric FISH probe (green). (D) Quantification of the TIF response as shown in (C) of POT1a^F/F^ MEFs before and after Cre treatment. Bar graphs represent averages from 3 experiments and SDs. p-values were derived from a two-tailed students t-test. p-values: *** ≤ 0.001; ** ≤ 0.01; * ≤ 0.05; n.s.: not significant. (E) Immunoblot for POT1a, POT1b and Stn1 in POT1a^F/F^ MEFs (with and without 96 h Cre treatment) expressing exogenous POT1b and/or Stn1 shRNA (indicated with +) or a scrambled sh (indicated with −). Asterisks indicate non-specific bands. Ctrl: non-specific band used as loading control. (F) Representative images of IF-FISH to monitor the DNA damage response at telomeres in the indicated Cre-treated (96 h) POT1a^F/F^ MEFs with and without POT1b and expressing Stn1sh or a scrambled control (ctrlsh). γH2AX foci that co-localize with telomeres (TIFs) were detected with IF with an α-γH2AX antibody (red) and FISH with a telomeric PNA probe (green). (G)Quantification of TIF response in the indicated POT1a^F/F^ MEFs (as in (F). Bar graphs represent averages from 3 independent experiments and SDs. P-values as in (D).

To confirm the findings with POT1bΔCST, we employed shRNA-mediated knockdown of Stn1. Consistent with the results with the POT1bΔCST mutant, partial knockdown of Stn1 restored the ability of overexpressed POT1b to repress ATR signaling at telomeres. In contrast, knockdown of Stn1 did not restore the ability of the endogenous POT1b to protect telomeres from ATR activation (Figure 3E-G). This is most likely due to the residual Stn1 interacting with endogenous POT1b, while the residual level of Stn1 is insufficient to bind all overexpressed POT1b. As CST is essential for cell viability, we are unable to test whether complete removal of Stn1 will restore ATR repression by endogenous POT1b.

## DISCUSSION

Because mammalian telomeres carry a 3’ protrusion of single-stranded DNA, ATR signaling has to be repressed throughout the cell cycle. Given the abundance of telomere ends in the nucleus, a system that fails at as few as 1% of the telomeres could lead to induction of cell cycle arrest and senescence or apoptosis. Our data clarify how this stringent ATR repression is achieved by the POT1 subunits of the telomeric shelterin complex.

### The mechanism of ATR repression at telomeres

The shelterin-tethered mutant version of RPA70 (Δ70) is remarkably proficient in repressing ATR signaling at telomeres, although not as effective as POT1a. The reduction in ATR signaling at telomeres containing Δ70-POT1aC suggest that the ability of POT1a to repress ATR signaling is primarily determined by its ability to bind to ssDNA telomeric DNA. Since the DNA binding domain of POT1a can be replaced the OB-folds of RPA70, we consider it unlikely that any other feature of POT1-DNA interaction beyond its binding to ssDNA is involved in the control of ATR signaling. The preference of POT1 for a TTAG-3’ telomere end appears to be largely irrelevant in this context. Indeed, TTAG-3’ ends only make up 40% of the telomere ends in human cells (Sfeir et al., 2005), yet all telomeres are shielded from the ATR pathway. Similarly, the ability of POT1 to protect G4 DNA from unfolding by RPA (Ray et al., 2014) is unlikely to be the primary determinant of ATR repression. Finally, although POT1 does not show a preference for its site next to the ds-ss junction in vitro (Palm et al., 2009), it could be argued that bound to shelterin, POT1 might have the ability to recognize the junction and prevent 9-1-1 loading. This hypothetical attribute is also rendered unlikely to be relevant to ATR repression based on the results with the shelterin-tethered chimeric RPA70 protein.

### The conundrum of POT1b and CST

The inability of POT1b to repress ATR signaling in S/G2 has presented a conundrum since POT 1a and POT 1b have identical TPP1- and DNA-binding features.

Furthermore, it can be inferred that POT1b resides at telomeres in S/G2 since it governs telomere end processing during this cell cycle stage. Our data indicate that the interaction of POT1b with the CST/Polα/primase complex prevents POT1b from acting as an ATR repressor. When the interaction of POT1b with CST is abrogated, POT1b is proficient in the repression of ATR in S/G2. This is an unexpected result since both POT1b and CST bind ss TTAGGG repeats.

### ATR repression at human telomeres

The single POT1 subunit in human shelterin has been implicated in the repression of ATR signaling (Hockemeyer et al., 2005; Denchi and de Lange, 2007). Interestingly, the recruitment of CST to human telomeres is mediated by TPP1, not POT1 (Wan et al., 2009). This arrangement may avoid the complication of CST interfering with the ability of POT1 to repress ATR signaling. Although human POT1 does play a role in how CST functions at telomeres (Takai et al., 2016; Pinzaru et al., 2016), we predict that CST recruitment by TPP1 does not interfere with the POT1-dependent exclusion of RPA from human telomeres.

## EXPERIMENTAL PROCEDURES

### Cell lines, expression constructs, and introduction of shRNAs

SV40 large T antigen (SV40LT) immortalized POT1a^F/F^ MEFs were reported previously (Hockemeyer et al., 2006). For expression of Myc-tagged proteins the following constructs were used: pWZL-Myc,(vector); Myc-POT1a (pWZL-Myc-POT1a); Myc-POT1aC (pWZL-Myc-POT1aC) containing POT1a aa 149-640 with an C-terminal Myc tag; Myc-Δ70-POT1aC (pWZL-Myc-Δ70-POT1aC) containing mouse RPA70 aa 191-623 fused to the N-terminus of POT1aC; Myc-ΔΔ70-POT1aC (pWZL-Myc-ΔΔ70-POT1aC) containing mouse RPA70 aa 191-331 fused to the N-terminus of POT1aC; Myc-Δ70 (pWZL-Myc-Δ70) containing RPA70 aa 191-623; Myc-SSB-POT1aC (pWZL-Myc-SSB-POT1aC) containing full length E. coli SSB fused to POT1aC; Myc-POT1b (pWZL-Myc-POT1b); and Myc-POT1bACST (pWZL-Myc-POT1bΔCST) (Wu et al., 2012). For each retroviral construct, 20 μg DNA was transfected into Phoenix packaging cells using CaPO_4_ co-precipitation. Medium was changed 12 and 24 h after transfection and the retroviral supernatant was used for 4 infections of the MEFs at 12 h intervals. Cells were subjected to selection in 135 μg/ml hygromycin for 3 days. To delete endogenous POT1a from POT1a^F/F^ MEFs, cells were subjected to three infections at 12 h intervals with pMMP Hit&Run Cre retrovirus derived from transfected Phoenix cells as previously described (Celli and de Lange, 2005). Time point 0 was set at 12 h after the first Hit&Run Cre infection. For Stn1 knockdown, the Stn1 shRNA (5’-GATCCTGTGTTTCTAGCCTTT-3’ (Wu et al., 2012)) in a pLKO.1 lentiviral vector (Openbiosystem) was produced in Phoenix-ECO cells as described above and introduced with 3 infections (6 h intervals). Infected cells were selected for 2.5 days in 2 μg/ml Puromycin.

### Co-immunoprecipitation

4×10^6^ HEK 293T cells were plated in a 15-cm dish 24 h prior to transfection using CaPO_4_ co-precipitation of 3 μg of pQE-StrepII-Rap1, pcDNA3.1-TRF2, pLPC-TIN2, pLPC-Myc-TPP1, pWZL-Myc-Δ70-POT1aC, pWZL-Myc-ΔΔ70-POT1aC as indicated. Medium was changed 8 h after transfection and cells were harvested 28 h later. Cells were resuspended in lysis buffer (60,000 cells/μl) (50 mM NaH_2_PO_4_ pH 8.0, 50 mM NaCl, 10 mM DTT, 0.5% Tween, 1x Complete protease inhibitors (Roche), 0.5 mM PMSF, and 8 μg/ml Avidin and incubated on a rotator at 4°C for 10 min. After centrifugation at 15,000 g at 4°C for 10 min, the supernatant was incubated with StrepII-beads (Qiagen; 1×10^6^ cells equivalent per μl beads) at 4°C for 1 h. Beads were washed 3 times in 500 μl lysis buffer and proteins were eluted with 20 μl of 2x Laemmli buffer (100 mM Tris-HCl pH 6.8, 200 mM dithiothreitol, 3% sodium dodecyl sulfate, 20% glycerol, 0.05% bromophenol blue). Samples were heated to 95°C for 5 min and analyzed by separation on 8% SDS-PAGE.

### Immunoblotting

Harvested MEFs (1×10^6^ cells) were resuspended in 100 μl 2x Laemmli buffer, incubated with 250 U Benzonase (Sigma-Aldrich) on ice for 10 min, denatured at 95°C for 5 min and separated on 8-16% SDS-PAGE (2×10^5^ cell equivalent per lane). After immunoblotting, the membranes were blocked in TBST with 2.5% not-fat dry milk and incubated with the following primary antibodies α-POT1a (#1221), α-Myc (9B11, Cell Signaling), α-TIN2 (#1447), α-Rap1 (#1253), α-RPA32 (A300-244A, Bethyl), α-RPA70 (A300-241A, Bethyl), α-POT1b (#1223), α-STN1 (sc-376450, Santa Cruz), α-tubulin (GTU88, Abcam) in 1% non-fat dry milk. Immunoblots for POT1a were performed using the renaturation protocol described previously (Loayza and de Lange, 2003). Secondary antibodies were horseradish peroxidase-conjugated donkey α-mouse IgG (GE Healthcare) or donkey α-rabbit IgG (GE Healthcare). Immunoblots were developed with enhanced chemiluminescence (Amersham).

### Protein expression and isolation

4×10^6^ HEK 293T cells were plated in a 15-cm dish 24 h prior to transfection using CaPO_4_ co-precipitation using 3 μg of pQE-StrepII-Myc-TPP1 (mouse), pLPC-Myc-POT1a, pLPC-Myc-Δ70-POT1aC, pLPC-Myc-ΔΔ70-POT1aC, and pLPC-Myc-SSB-POT1aC as indicated. Medium was changed 8 h after transfection and cells were harvested 28 h later. Subsequently, cells were pelleted, washed twice in PBS, resuspended in lysis buffer (60,000 cells/μl) (50 mM NaH_2_PO_4_ pH 8.0, 50 mM LiCl, 10 mM DTT, 0.5% Tween, 1x Complete protease inhibitors, 0.5 mM PMSF, and 8 μg/ml Avidin) and incubated at 4°C for 10 min on a rotator. After centrifugation at 15,000 <u>g</u> at 4°C for 10 min, the supernatant was incubated with Strep-tactin beads (Qiagen; 1×10^6^ cells equivalent per μl beads) and 250 U Benzonase (Sigma-Aldrich) for 1 h at 4°C on a rotator. Beads were washed 3 times in lysis buffer and eluted twice in Elution buffer (50 mM NaH_2_PO_4_ pH 8.0, 50 mM LiCl, 1 mM DTT, 0.1% Tween, 10 mM Biotin). Samples were aliquoted, snap-frozen and stored at −80°C.

### Electrophoretic mobility shift assay

Labeling reactions and gel-shift assays were performed as described previously (Palm et al., 2009) with minor modifications. All oligonucleotides were obtained from Sigma. The probe sequences were as follows: Telo34 (5’-TTAGGGTTAGGGTTAGGGTTAGGGTTAGGGTTAG-3’); Scr35 (5’-ATGCGACTCGAGCTAGATGATGTCTTCTGCAATCA-3’). Gel-shift reactions were performed in 10 μl reaction buffer (50 mM Hepes-KOH pH 8.0, 50 mM LiCl, 0.1 mM DTT, 50 ng/μl ß-Casein) with 0.2 nM Polynucleotide kinase end-labelled DNA probes. Proteins were added last and the mixture was incubated for 10 min on ice. Electrophoresis was performed in 0.8% agarose gels run in 0.5x Tris-borate/EDTA pH 8.3. Gels were run for 45 min at 100 V at room temperature, fixed in 20% methanol/10% acetic acid, dried on Whatman DE81 paper at 80°C with vacuum and exposed to phosphorimager screens. The binding fractions were calculated taking all protein-DNA complexes into account [binding fraction (%)] = [proteins-DNA complex]/[free DNA probes] to derive apparent relative K_d_ values.

### IF, IF-FISH, and TIF analysis

Cells grown on Poly-lysine covered coverslips were harvested at time point 96 h after Cre to analyze protein localizations and TIF response. Coverslips were incubated on ice, rinsed once with PBS containing Mg^2+^ and Ca^2+^ and soluble proteins were pre-extracted with ice cold Triton-X 100 buffer (20 mM Hepes-KOH pH 7.9, 50 mM NaCl, 3 mM MgCl_2_, 0.5% Triton X-100, 300 mM Sucrose) for 30 sec on ice. Coverslips were rinsed twice with cold PBS containing Mg^2+^ and Ca^2+^, transferred to room temperature and fixed in 3% paraformaldehyde with 2% sucrose for 10 min. Coverslips were washed twice in PBS at room temperature for 5 min. IF was carried out as previously described (Takai et al., 2003; Celli and de Lange, 2005; Dimitrova and de Lange, 2006) with the following primary antibodies: α-Myc (9B11, Cell Signaling), α-γH2AX (#05-636, Millipore), α-53BP1 (ab175933, Abcam). For IF-FISH staining, after the secondary antibody incubation and wash step, cells were fixed again with 2% paraformaldehyde for 5 min; dehydrated in 70%, 95%, and 100% ethanol for 5 min each; and allowed to air dry. Hybridizing solution (10 mM Tris-HCl pH 7.2, 70% formamide, 1 mg/ml blocking reagent [Roche], containing 100 nM of a fluorescent PNA probe (488-OO-[CCCTAA]3 [PNA Bio Inc.]) was added to each coverslip, and the DNA was denatured by heating for 10 min at 80°C on a heat block. After 2 h of incubation at room temperature in the dark, cells were washed twice with washing solution (70% formamide, 10 mM Tris-HCl pH 7.2) for 15 min each and with PBS three times for 5 min each. DAPI (0.1 μg/ml) was added to the second PBS wash. Coverslips were sealed onto glass slides with embedding medium (ProLong Gold anti-fade reagent, Invitrogen). Digital images were captured on a Zeiss Axioplan II microscope with a Hamamatsu C4742-95 camera using Volocity software.

### S phase index

Before harvest, cells seeded on coverslips were incubated for 1.5 h in medium containing 10 μM BrDU and subsequently fixed with 2% PFA at room temperature for 10 min. Following an incubation in NP-40 buffer (0.5% NP-40 in PBS) at room temperature for 10 min, coverslips were washed 3x in PBS for 5 min and then incubated in 4N HCl at room temperature for 10 min. Next, the coverslips were washed 3x in PBS for 5 min and incubated with Blocking solution (PBS, 0.1% BSA, 3% Serum Donkey, 0.1% Triton-X 100, 1 mM EDTA pH 8.0) at room temperature for 45 min before addition of α-BrdU antibody (B35130, Invitrogen). Cells were incubated with the antibody at 4°C for overnight. Coverslips were washed 3 times in PBS for 5 min and incubated with secondary antibody at room temperature for 45 min. Coverslips were washed 3 times in PBS for 5 min, with the second PBS-wash containing DAPI (0.15 μg/ml) to stain for DNA. Coverslips were sealed and processed as described for IF above.

### Telomeric overhang analysis

Mouse telomeric DNA was analyzed on CHEF gels as described previously (Wu et al., 2012).

## ACKNOWLEDGEMENT

We thank Hiro Takai and Melissa Pamula for executing initial experiments and members of the de Lange lab for helpful discussion. KK was supported by an EMBO long-term fellowship (EMBO ALTF-1044-2011) and funds from the *Women & Science* Fellowship Program at The Rockefeller University. This work was supported by a grant from the NIH (AG016642) to TdL. TdL is an American Cancer Society Rose Zarucki Trust Research Professor.

## CONFLICT OF INTEREST

The authors declare that they have no conflicts of interest with the contents of this article.

## SUPPORTING FIGURES

**Figure S1.**
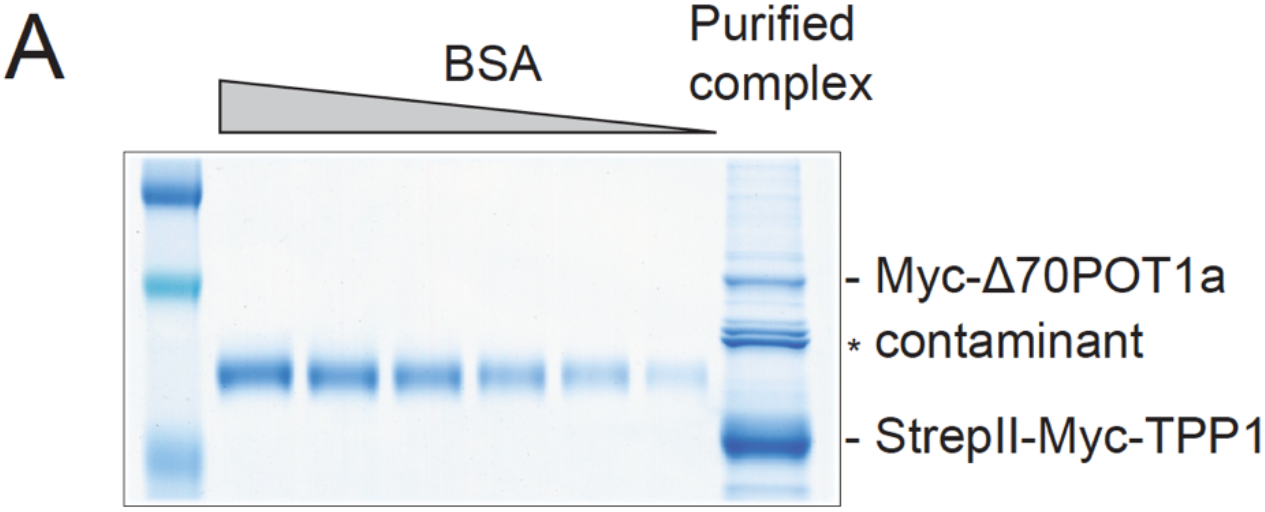
Isolation of Δ70-POT1aC. TPP1 and Δ70-POT1aC were co-expressed in HEK293T cells and isolated using the StrepII-tag of TPP1. The Coomassie gel shows a BSA standard to determine the protein concentration of Δ70-POT1aC to calculate the relative K_d_s in Figure 1G.

**Figure S2.**
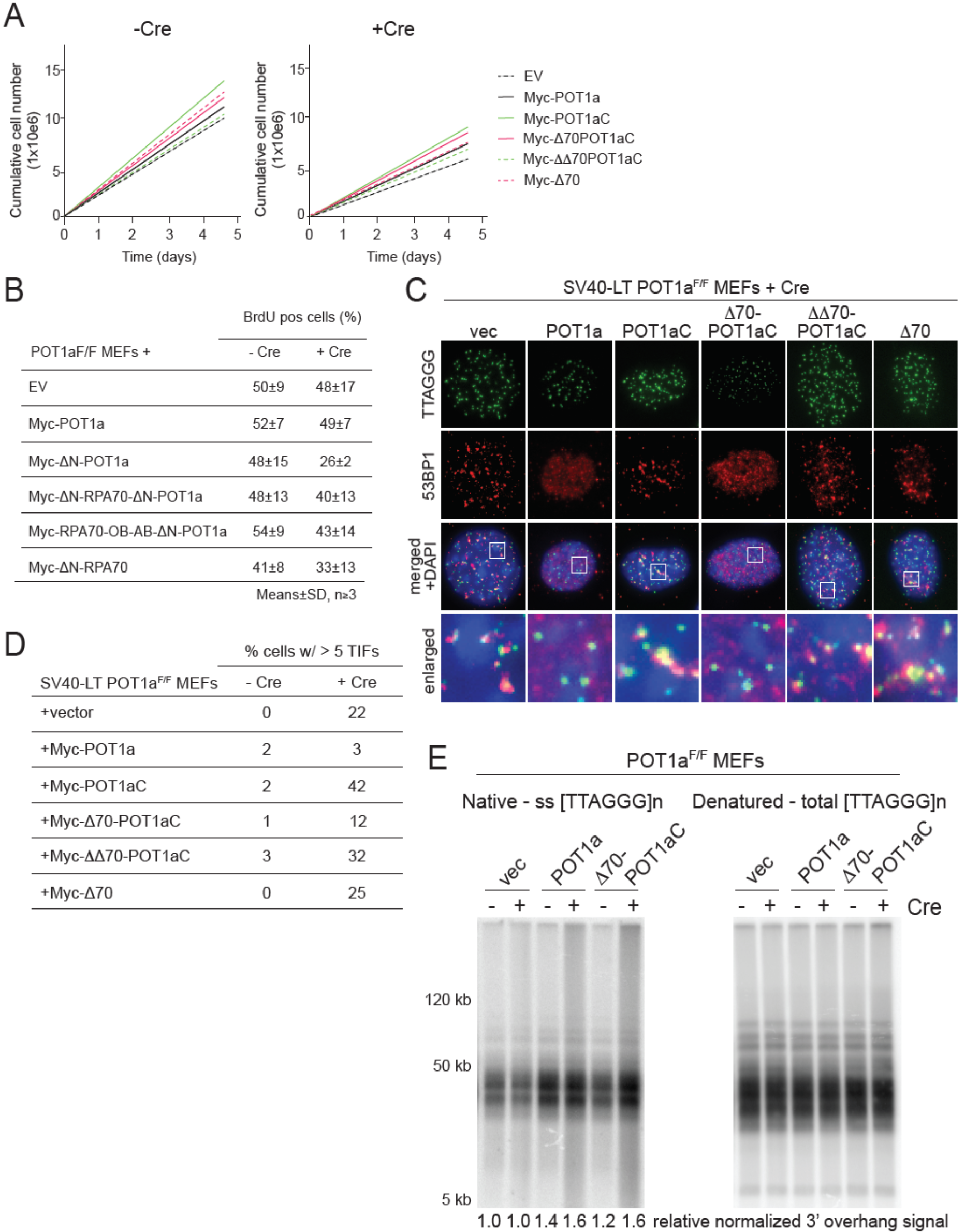
Characterization of RPA-POT1aC expressing cell lines. (A) Proliferation rate of the different cell lines used. (B) S phase indices of POT1a^F/F^ MEFs with and without Cre treatment (96 h) expressing the indicated proteins. Average of 3 independent experiments and SDs. (C) Representative IF images of the DNA damage response at telomeres of Cre-treated (96 h) POT1a^F/F^ MEFs expressing the indicated proteins. 53BP1 foci that co-localize with telomeres (TIFs) were detected with α-53BP1 antibody (red) combined with FISH for telomeric DNA (green). (D) Quantification of the TIF response (53BP1 foci at telomeres) in POT1a^F/F^ MEFs expressing the indicated proteins with and without Cre treatment (96 h) as shown in (C).
Quantitative analysis of the ss TTAGGG repeats at telomeres of POT1a^F/F^ MEFs with and without Cre treatment expressing the indicated proteins (96 h). Numbers below the native gel (left) represent the relative 3’ overhang signal normalized to the total TTAGGG repeat signal in the same lane.

**Figure S3.**
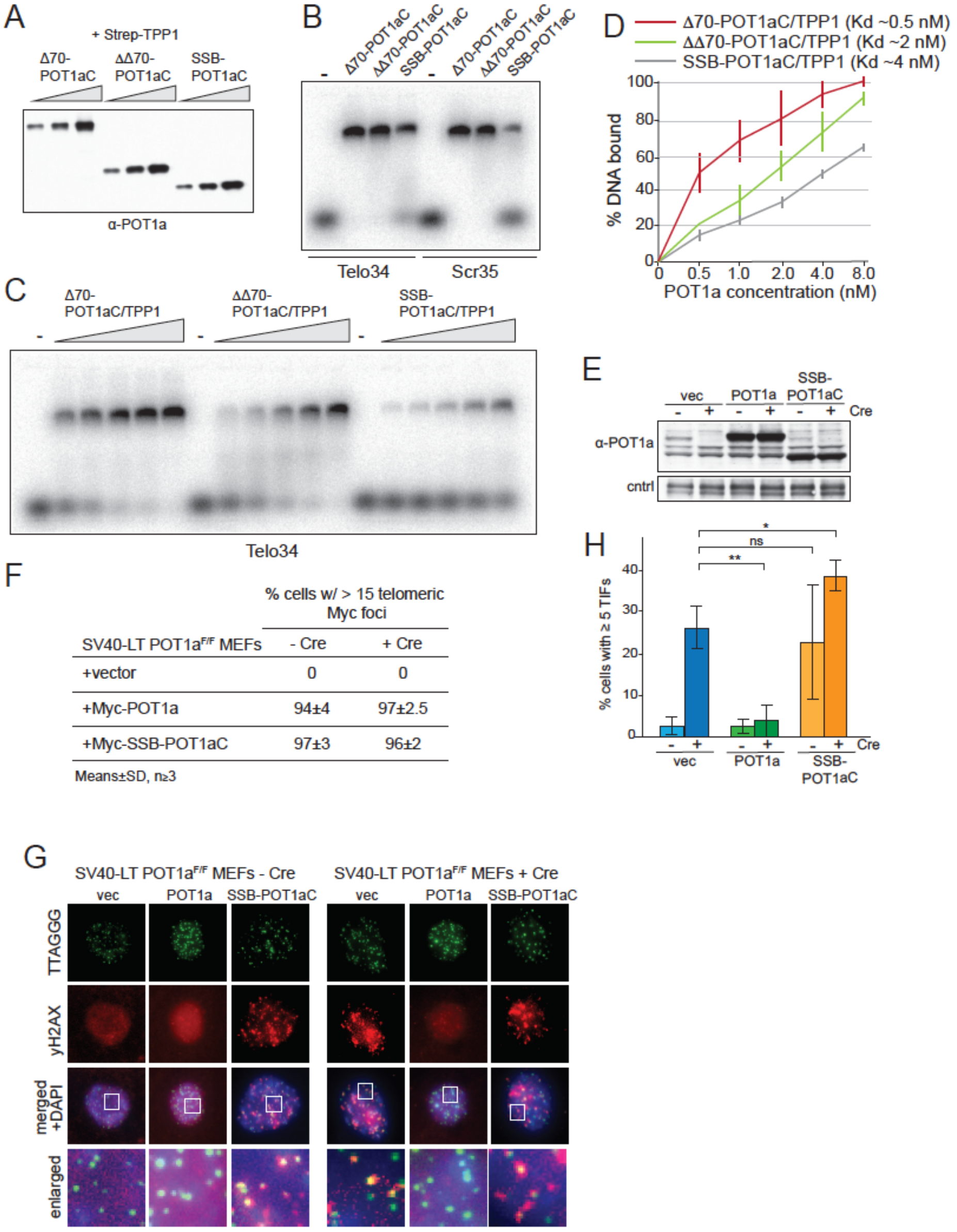
Shelterin-tethered SSB does not repress ATR. (A) Immunoblot to determine the relative protein concentration of Strep-tagged TPP1 in complex wtih either the indicated RPA-fusion proteins or SSB-POT1C isolated from transfected 293T cells. (B) EMSA showing binding of TPP1/Δ70-POT1 aC, TPP1/ΔΔ70-POT1aC, and TPP1/SSB-POT1aC to telomeric and non-telomeric single-stranded DNAs. (C) Representative EMSA to determine the apparent relative K_d_ of TPP1/Δ70-POT1aC, TPP1/ΔΔ70-POT1 aC, and TPP1/SSB-POT1aC for Telo34. (D) Quantification of the affinity of TPP1/Δ70-POT1aC, TPP1/ΔΔ70-POT1aC and TPP1/SSB-POT1aC as shown in (C) from 3 independent experiments. Error bars represent SDs. (E) Immunoblot for POT1a in SV40-LT immortalized POT1a^F/F^ MEFs expressing the indicated proteins before and after Cre treatment (96 h). The endogenous POT1a is lost upon Cre treatment and the introduced proteins are overexpressed compared to endogenous POT1a. (F) Quantification of the telomeric localization of the indicated proteins in POT1a^F/F^ MEFs with and without Cre treatment (96 h). Averages of 3 independent experiments and SDs. (G) Representative IF images of the DNA damage response at telomeres of Cre-treated (96 h) POT1a^F/F^ cells expressing the indicated proteins. γH2AX foci that co-localize with telomeres (TIFs) were detected with an α-γH2AX antibody (red) combined with FISH for telomeric DNA (green). (H) Quantification of the TIF response as shown in (G) in the indicated POT1a^F/F^ cells with and without Cre treatment. Data represent averages from 3 independent experiments. P values as in Figure 2.

